# MAGMA: inference of sparse microbial association networks

**DOI:** 10.1101/538579

**Authors:** Arnaud Cougoul, Xavier Bailly, Ernst C. Wit

## Abstract

Microorganisms often live in symbiotic relationship with their environment and they play a central role in many biological processes. They form a complex system of interacting species. Within the gut microbiota these interaction patterns have been shown to be involved in obesity, diabetes and mental disease. Understanding the mechanisms that govern this ecosystem is therefore an important scientific challenge. Recently, the acquisition of large samples of microbiota data through metabarcoding or metagenomics has become easier.

Until now correlation-based network analysis and graphical modelling have been used to identify the putative interaction networks formed by the species of microorganisms, but these methods do not take into account all features of microbiota data. Indeed, correlation-based network cannot distinguish between direct and indirect correlations and simple graphical models cannot include covariates as environmental factors that shape the microbiota abundance. Furthermore, the compositional nature of the microbiota data is often ignored or existing normalizations are often based on log-transformations, which is somewhat arbitrary and therefore affects the results in unknown ways.

We have developed a novel method, called MAGMA, for detecting interactions between microbiota that takes into account the noisy structure of the microbiota data, involving an excess of zero counts, overdispersion, compositionality and possible covariate inclusion. The method is based on Copula Gaussian graphical models whereby we model the marginals with zero-inflated negative binomial generalized linear models. The inference is based on an efficient median imputation procedure combined with the graphical lasso.

We show that our method beats all existing methods in recovering microbial association networks in an extensive simulation study. Moreover, the analysis of two 16S microbial data studies with our method reveals interesting new biology.

MAGMA is implemented as an R-package and is freely available at https://gitlab.com/arcgl/rmagma, which also includes the scripts used to prepare the material in this paper.

## 1 Introduction

Microbiota are ubiquitous and play a central role in biological processes [1]. High-throughput sequencing allows to study the composition, structure and diversity of complex microbial communities. In the wake of technological development, there has been during the last years a multiplication of projects querying the structure and properties of specific microbiota. Among others, some large projects targeted the human microbiome, e.g., the MetaHIT project [2, 3] and the HMP project [4, 5, 6], planktonic and coral ecosystems of the different oceans (TARA Oceans) project [7, 8], or the earth’s multiscale microbial diversity (EMP) project [9, 10].

Microbiota are by nature complex systems of interconnected taxa. Interactions among microbes are an important factor that shape the structure and properties of microbiota. From an ecological point of view, interactions appear to structure [11], stabilize [12] and regulate the diversity [13] of microbial communities. In the biomedical field the dysbiosis of the human gut microbiota is associated with multiple pathologies such as obesity [14], diabetes [15] and mental illness [16]. Metagenomics opens a field of exploration of potential associations between the microbiome and several complex diseases [17]. Global modifications of a microbiota can also have implications for the dynamics of a bacteria of particular interest. In epidemiology the infection of a host by a pathogen can be facilitated by some microbial species through various interaction processes [18]. Conversely, some microbial species may have antagonistic interactions with pathogens that could be used in biological control [19, 20].

Identifying potential microbial interactions from metagenomic data is therefore a topical scientific challenge. Methodological developments are needed to improve this identification, taking into account the noisy and stochastic structure of the genomic measurement process of the microbiota.

### 1.1 Metagenomic data characteristics

Metagenomic data from 16S rRNA sequencing consists of sequencing reads originating from thousands of different bacterial groups obtained from hundreds to thousands of samples [21]. In order to reflect the microbial composition and the relative frequency of each bacterial group among samples, sequencing reads are clustered in Operating Taxonomic Units (OTU) [22], e.g., bacterial species. The number of OTUs considered depends both on the studied microbial community and the criteria used to cluster sequences in OTUs.

Metagenomic read counts are sparse and overdispersed. Most of OTUs are rare and occur in only a few samples. The sequencing read data therefore have a large amount of zeros [23], which often in naive analyses causes spurious associations [24]. Zero-inflated (ZI) distributions thus appear to be the most appropriate to model OTU abundances. Furthermore, the abundance of an OTU, defined as the number of reads assigned to this OTU, does not follow a usual count distribution such as a zero-inflated Poisson distribution as typically overdispersion is observed when the OTU is present. It has been shown that ZI negative binomial or ZI lognormal provide a good fit [25, 26].

Another aspect to take into consideration is the sequencing depth of a sample, which is defined as the sum of all OTU read counts in a sample. Sequencing depths are unequal among samples due to experimental effects [27]. From this perspective, metagenomic sequencing read data should be considered compositional in nature [28, 29]. For a given sample, each OTU read abundance “depends” on the other OTU reads through the sequencing depth. various ways have been suggested for taking the sequencing depth into account when analyzing the observed read counts for an OTU. A common way is to circumvent the compositional nature of the data and to make OTUs comparable by transforming and normalizing the OTU table before further analysis. Main methods are rarefying, scaling and log ratio transformation, but all have problematic aspects [25, 26, 30]. A typical scaling transformation method is to divide by the marginal sum of the sample, i.e., the sequencing depth. This method leads to spurious negative associations [31]. The centered log ratio (clr) transformation [32] and relative log expression [33] are most commonly used to process compositional sequencing data. The microbial data is mainly composed of zeros and the log cannot be applied without replacing zero values by a pseudo-count and is therefore not ideal [26, 34].

The diversity of microbiota among samples furthermore depends on factors that are known to structure the distribution of microbes such as environmental conditions, spatial and temporal scales. For instance, age, genetics, environment and diet are all factors that affect the human gut microbiota [35]. Seasonal changes in the microbiota of wild mice have also been observed [36]. These factors should be considered as much as possible in the analysis by integrating them as covariates in the model to separate biological interactions from the effects of structuring covariates.

### 1.2 Inference of microbial associations networks

In this paper, we take the perspective to consider microbial communities as a network of microbial species (or OTUs) that interact with each other. These networks are formalized by graphs consisting of vertices representing OTUs and edges representing statistical dependencies, i.e. associations, between OTUs. Network analysis is the most common approach to explore potential microbial interactions at the microbiota scale. There are two main ways to infer a microbial network: correlation-based networks and graphical models [37].

On the one hand, correlation-based networks are graphs obtained from computing and thresholding all pairwise association measures. A large number of association measures have been used in this framework, such as correlation (e.g., Pearson, Spearman), similarity (e.g., mutual information), or dissimilarity (e.g., Kullback-Leibler) measures [38]. These methodologies rely on pairwise associations between occurrences or abundances of bacterial OTUs among the microbiota. A permutation and bootstrap approach can be used to improve the robustness of the infered network [39]. The main disadvantage of pairwise association methods is that they are unable to distinguish between direct and indirect associations, thereby often ending up with dense network that give little insight in the underlying functional relations.

On the other hand, graphical models have minimal bias and better power [37, 40]. Graphical models are graphs that satisfy the Markov properties, which means that links represent conditional dependencies. In the multivariate Gaussian case, conditional dependence is equivalent to a non-zero partial correlation. In a such framework, the conditional dependencies can be read off from the inverse correlation matrix, called the precision matrix. Inference of Gaussian graphical models can be performed by neighborhood selection [41] or by lasso regularization [42]

The two main methods used for exploration of microbial interactions are SPIEC-EASI [40] and SparCC [43]. SparCC estimates linear Pearson correlations between the log-transformed components. The algorithm works by iteratively calculating a “basis correlation” under the assumption that the majority of pairs do not correlate [43]. SPIEC-EASI normalizes the data with the *clr* transformation before applying the classical framework of Gaussian graphical models described below. Both methods use a pseudo-counts to avoid zeros and can not take into account potential covariates.

Current network inference methods such as SPIEC-EASI and SparCC do not fully consider the structure of metagenomic data involving sparsity, overdispersion, compositionality or covariate inclusion. We therefore propose a novel inference framework involving copula Gaussian graphical models [44]. This model provides a general and integrative framework for network inference. We called our method MAGMA for **M**icrobial **A**ssociation **G**raphical **M**odel **A**nalysis. MAGMA allows to take into account all aspects of the data, while relying on the well-known properties of a latent Gaussian graphical model. We implemented our method in R and provide a package called `rMAGMA` available on a Git repository at https://gitlab.com/arcgl/rmagma.

## 2 Materials and Methods

Here we present an original way of integrating metagenomic data for the exploration of microbe-microbe interactions. We propose a copula Gaussian graphical model combined with GLM marginal distributions. Although full likelihood inference is possible, our MAGMA approximation is based on the estimation of the latent data by the median of possible values. This mapping makes it possible to manage the excess of zeros, overdispersion, the compositional nature of the data and the inclusion of covariates.

### 2.1 Model

In the classical Gaussian graphical model (GGM), we consider a centred multivariate normal *Z* with a correlation matrix Θ^−1^,

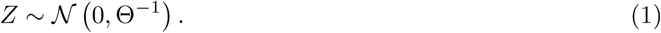

Computing the precision matrix Θ gives informations about partial correlations between elements of *Z* [45]. Under the multivariate Gaussian assumption, the partial correlation *ρ*_*ij*_ between *i* and *j* is given by:

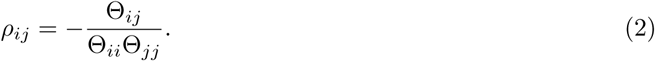

Non-zero elements in the precision matrix Θ correspond to the conditional dependencies and edges in the conditional dependence graph.

The observed metagenomic count data, unfortunately, do not follow a normal distribution. Microbiota data are represented by a matrix *Y* of *n*×*p* dimension, where *n* is the number of samples and *p* is the number of OTUs. We assume that the joint distribution of observed variables *Y* can be transformed from a latent multivariate normal variable *Z*. The copula Gaussian graphical model defines the marginal transformations [44],

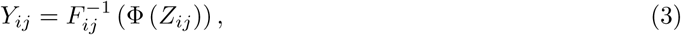

where Φ is the cumulative distribution function (cdf) of the standard normal distribution and 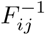 is the inverse cdf of microbiota count *Y*_*ij*_ for the *j*^*th*^ OTU and for sample *i*.

The *F*_*ij*_ function is generally estimated by the empirical cdf 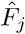 [46, 47], but this is not appropriate here as *F*_*ij*_ will certainly depend on the sequencing depth of sample *i* and therefore cannot be constant across samples. Instead, we assume that OTU read abundances are distributed according to a zero-inflated negative binomial (ZINB). We introduce the original mapping function:

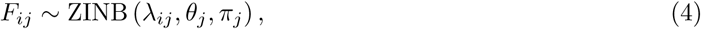

where *λ*_*ij*_ is the mean of the negative binomial part for sample *i* and species *j, θ*_*j*_ is the dispersion parameter and *π*_*j*_ is the probability of the structural zeros. The mean *λ*_*ij*_ is defined by the equation:

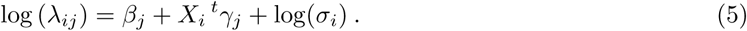

*β*_*j*_ is modelling the mean of species *j, γ*_*j*_ is the effect of covariates *X* on species *j* and *σ*_*i*_ is the library size or sequencing depth for sample *i*.

With this parametric mapping function, we can model the high proportion of zeros in data by the use of a zero-inflated distribution. We model overdispersion by the negative binomial distribution. We model sequencing depth to take into account compositionality by an offset. And we also model the effect of covariates, either qualitative or quantitative, on the mean of microbial abundance.

### 2.2 MAGMA inference

Full likelihood inference of the above model is involved. We propose here a computational approximation of the maximum likelihood. If *Y* were continuous data, then observed variables could be projected into the latent space by the inverse mapping,

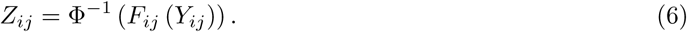

But since *Y*_*ij*_ are discrete count data, 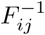 is not injective and the projection in the latent space is not unique. *F*_*ij*_ is a step function and *Z*_*ij*_ can take all the values in the interval [Φ^−1^ (*F*_*ij*_ (*Y*_*ij*_ − 1)), Φ^−1^ (*F*_*ij*_ (*Y*_*ij*_))].

To approximate the copula Gaussian graphical model, the nonparanormal normal score approach [48] takes the right bound value 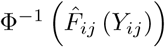 and winsorizes the data for the highest observed values to avoid infinite values. The nonparanormal SKEPTIC transformations [49] use the asymptotic relationships between the Pearson correlation and the Spearman or Kendall rank correlations. Instead, we propose to transform the count data using the median point of the Z distribution of reachable values,

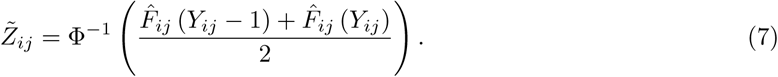

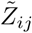 thus defined is the median of the normal distribution between 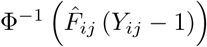 and 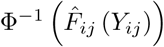. With this estimation, we do not need to winsorized the data nor rely on dubious asymptotic relationships that certainly do not hold.

For estimating the library size *σ*_*i*_ of sample *i* in (5), the sample sequencing depth ignores the fact that different biological samples may express different 16S RNA repertoires [50]. We estimate the library size using the geometric mean of pairwise ratios (GMPR) [51]. GMPR is specifically intended for compositional zero-inflated data as the microbiome sequencing data. For each pair of samples *i* and *i*′, the median of count ratios of nonzero counts is computed,

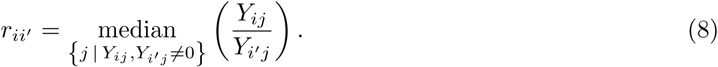

The ratio *r*_*ii′*_ represents how much, on average, the OTU read counts of sample *i* are above or below those of a sample *i*′. If *r*_*ii′*_ = 2, the OTU of the sample *i* will have on average 2 times more read counts than those of sample *i*′. To estimate the library size factor of a sample *i*, we then compute the geometric mean of all the ratios *r*_*ii′*_ involving the sample *i*. This is the average difference between the abundance of an OTU found in sample *i* and its abundance in the other samples,

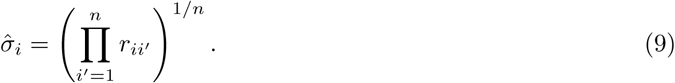

The GLM (5) is then estimated with off-sets 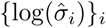, which then allows us to calculate the quasi-normal data 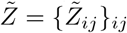 according to (7).

Finally, we propose to infer the association network from the transformed data 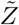 of the observed variable *Y*. In this way, we approximately infer the copula Gaussian graphical model, taking into account the characteristics of the microbial data to infer relevant associations between OTUs. We use graphical lasso (glasso) inference [42] from the `R huge` package to estimate a sparse precision matrix. In the sparse estimation of the precision matrix Θ, the problem is to maximize the penalized log likelihood

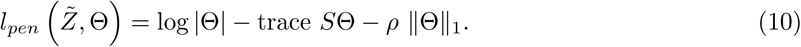

*S* denotes the empirical covariance of the 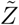 transformed data matrix, |Θ|_1_ is the *L*_1_ norm and 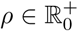 is a sequence of non-negative penalty parameters.

Penalized inference of graphical models results in a collection of OTU networks associated with the estimated precision matrix 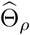 for different values of *ρ*. In order to infer the most parsimonious network given the available data, one need to weigh the fit of the data relative to the complexity of the data [52, 53]. To select the penalty parameter *ρ*, we consider three approaches: rotation information criterion (ric) [54], stability approach for regulation selection (stars) [55] and extended Bayesian information criterion (ebic) [56]. All these approaches are encoded in the R package `huge` used by MAGMA.

In summary, MAGMA inference comprises of the following steps:

1. Adjust the marginal OTU abundances to ZINB distributions according to equations (4) and (5).
2. Approximate the latent data *Z* according to equation (7).
3. Estimate a sparse precision matrix 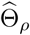 according to equation (10).
4. Select the penalty *ρ*^*^ that best balances fit and complexity via ric/stars/ebic.
5. Identify the OTU network from non-zero elements of 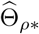.

## 3 Results and Discussion

We studied the efficiency of the MAGMA tool to infer a network of microbial associations. With this aim, we first analyzed the behavior of MAGMA on simulated data in section 3.1. We measured the quality of network inference under different conditions and compared MAGMA with other network approaches. In section 3.2 we applied MAGMA to data from the Human Microbiome Project.

### 3.1 Simulation study

In this section, we first describe how we generate simulation data. We then studied six different aspects of the MAGMA model with respect to this simulated data: (i) its consistency, i.e., whether it converges to the true network with increasing number of samples *n*, (ii) its robustness, i.e., whether it is able to deal with deviations from ZINB read counts, (iii) its ability to infer the network with varying interaction strengths, (iv) how its ability to reconstruct the network depends on different network topologies, (v) its ability to account for confouding by integrating a covariates, and finally, (vi) we compared MAGMA with existing tools for the inference of microbial association networks. The ability of the procedure to recover the simulated microbial network was measured via the area under the ROC curve (AUC) along the *ρ*-path of the inferred networks.

#### 3.1.1 Generation of realistic data sets

To measure the performance of network inference tools, we should simulate datasets of known structure and tried to recover the associations that we simulated. SPIEC-EASI [40] proposes a simulation procedure, however it is unable to reproduce variations in sequencing depth, which is considered an essential feature [57]. Our procedure first generates an association network *G* with *d* vertices and *e* edges (Figure 1). The topology of the generated network can be selected to be either band, block, cluster, hub, random or scale free as defined in [40]. We associate the simulated graph with an inverse correlation matrix fixing the condition number of the matrix *k* regulating the strength of correlations. We then generate multivariate normal data with the obtained correlation structure.

**Figure 1:**
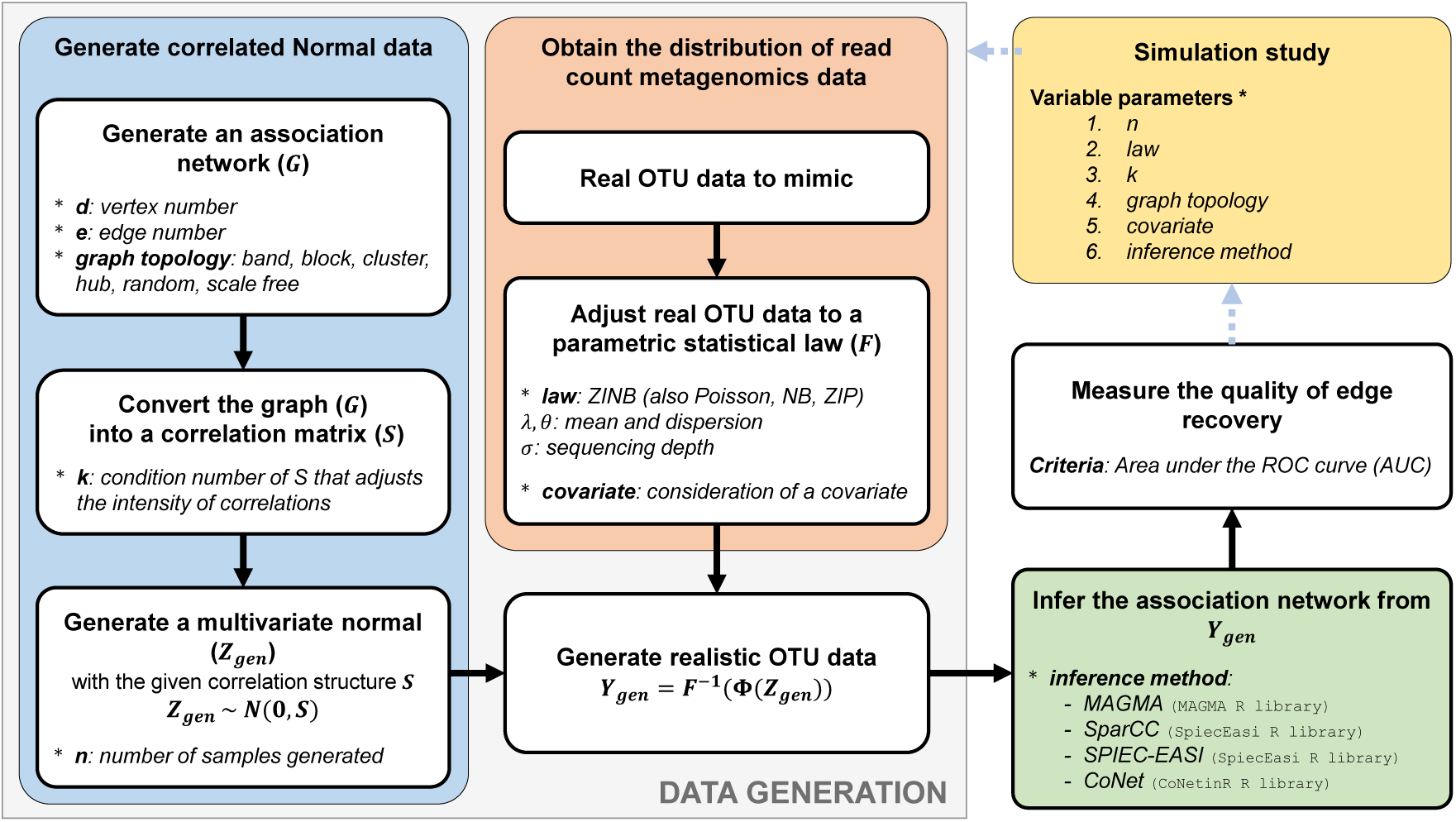
Workflow of the generation of realistic data for the inference benchmarking.

Then we need to transform the latent normal data into the observed read count data. To mimic the structure of real data, we relied on the 16S data of the microbiome of Puerto Rico honey bees obtained by MG Dominguez-Bello [58, study ID 1064]. We filtered the data, keeping the 80 OTUs with a prevalence greater than 15% and 286 samples with a sequencing depth greater than 100 reads. The average sequencing depth was 19,000. The data has been fitted according to some parametric distribution, e.g., the ZINB used in our network inference, but also other distributions: Poisson, zero-inflated Poisson and negative binomial. Using the copula transformation, we project the multivariate normal data into read counts using the selected marginal distributions combined with the logarithmic link function involving covariates and an offset.

#### 3.1.2 Effect of the sample size

Data were simulated with different number of samples *n*. We then inferred the association network and measured the quality of edge recovery as shown in Figure 2A. As the number of samples increases, the AUC increases and tends to one. Asymptotically, the method correctly recovers all the simulated links. The approximation of the copula Gaussian graphical model made by MAGMA allowed to recover the network with hundreds of samples. With 200 simulated OTUs, 200 to 300 of samples are sufficient recover almost the entire network correctly.

**Figure 2:**
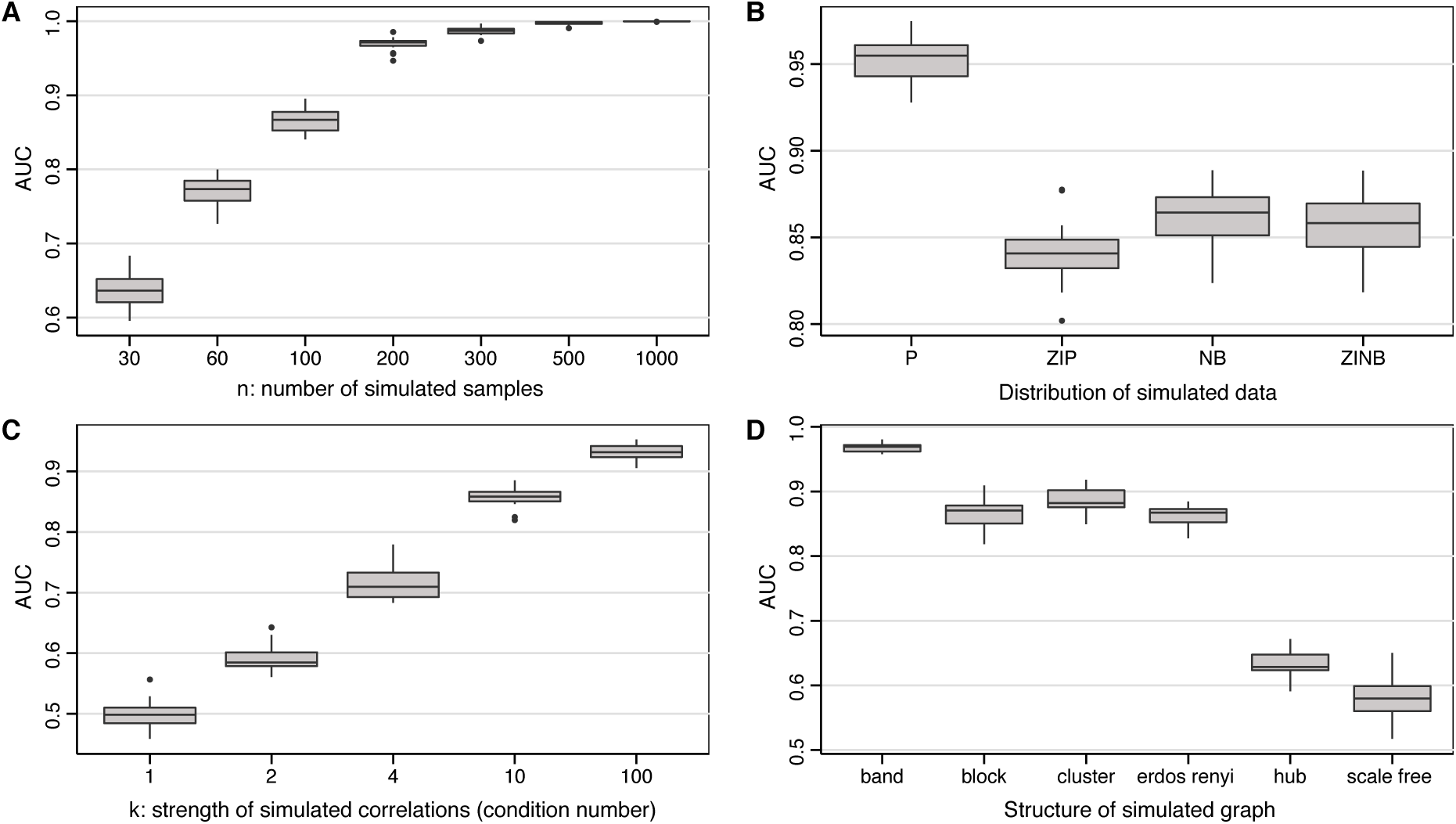
Effect of varying parameters on the quality of network inference. Boxplots of the AUC criterion according to: (**A**) the number of simulated samples *n* varying from 30 to 1000; (**B**) the distributions of the simulated data (Poisson, Zero-inflated Poisson, Negative Binomial and Zero-inflated Negative Binomial); (**C**) the condition number of the simulated correlation matrix *k* varying from 1 to 100; (**D**) the structure of simulated graph (band, block, cluster, hub, random and scale free). If they did not vary, the parameters were fixed at: *n* = 100, *k* = 10, random graph structure (Erdos-Renyi), marginal count data simulated according to a ZINB. We considered 20 simulation iterations for networks of size 200 with an average degree of 2.

#### 3.1.3 Effect of the distribution of read counts

To measure the flexibility of our method to model misspecification, we varied the distribution of the simulated data. The results are shown in Figure 2B. Zero-inflated Poisson, negative binomial and zero-inflated negative binomial all performed roughly similar, suggesting that MAGMA is quite robust to model misspecification, as it assumes underlying ZINB data. It is striking that dependence networks with underlying Poisson distributed read count data were able to be reconstructed significantly better (average AUC > 0.95), suggesting that the zero-inflation and, particularly, over-dispersion makes network reconstruction more difficult (average AUC ≈ 0.85).

#### 3.1.4 Effect of the strength of partial correlations

The strength of the simulated correlations was modelled by the condition number of correlation matrix of the simulated data. A low condition number corresponds to small values of the coefficients of the correlation matrix and this will produce weak links. The results are shown in Figure 2C. With a condition number of 1, its lowest possible value, the correlations have no strength and the results obtained were the same as a random draw with an average AUC of 0.5. The AUC increases rapidly with an increasing condition number. For *k* = 4, we found an AUC at 0.72 and for *k* = 10 the AUC was already at 0.85.

#### 3.1.5 Effect of network topologies

As Figure 2D shows, network topology has, perhaps surprisingly, a significant impact on network reconstruction quality. Simulations were done for different kind of graph structures. The band graph has the best reconstruction properties (average AUC > 0.95). On the other end, the recovery of a scale free or a hub network was difficult (average AUC ∈ (0.55, 0.65)). It seems that high-degree nodes pose a problem with network inference. This is a common issue also with other methods [40]. The reason why the band topology can be easily reconstructed may be because it has the lowest maximum node degree of all topologies. The results for the block, cluster and random networks were good with an AUC above 0.85.

#### 3.1.6 Consideration of a covariate

In order to check the capacity of our method to account for confounding in the dependence network, we used MAGMA with the inclusion of a quantitative covariate. We generated read count data by adding a covariate effect with different levels of strength. The coefficients of the covariate, *γ*_*j*_ in (5), for all OTUs were sampled from a normal distribution with variance equal to 0, 1, 2 or 4. The mean of *γ* is taken to be zero, as it is just an offset, confounded with the sampling effort. The values of the unit specif covariate are sampled from a standard normal distribution.

We compare the effect of including and ignoring the covariate effect across different levels of confounding. As Figure 3 shows, when there is in fact no confounding, using MAGMA containing an irrelevant covariate does not result in more errors than using MAGMA without the covariate. The addition of an irrelevant covariate effect to the method does not have a negative impact on the AUC. As the strength of the confounding increases, MAGMA that accounts for this confounding has an increasing advantage in recovering the network structure over applying MAGMA that ignores the covariate. We conclude that careful modelling of the read count distribution *F*_*ij*_ is particularly relevant: the inference quality of the association network relative to the agnostic MAGMA increases when the covariate effect is gets stronger.

**Figure 3:**
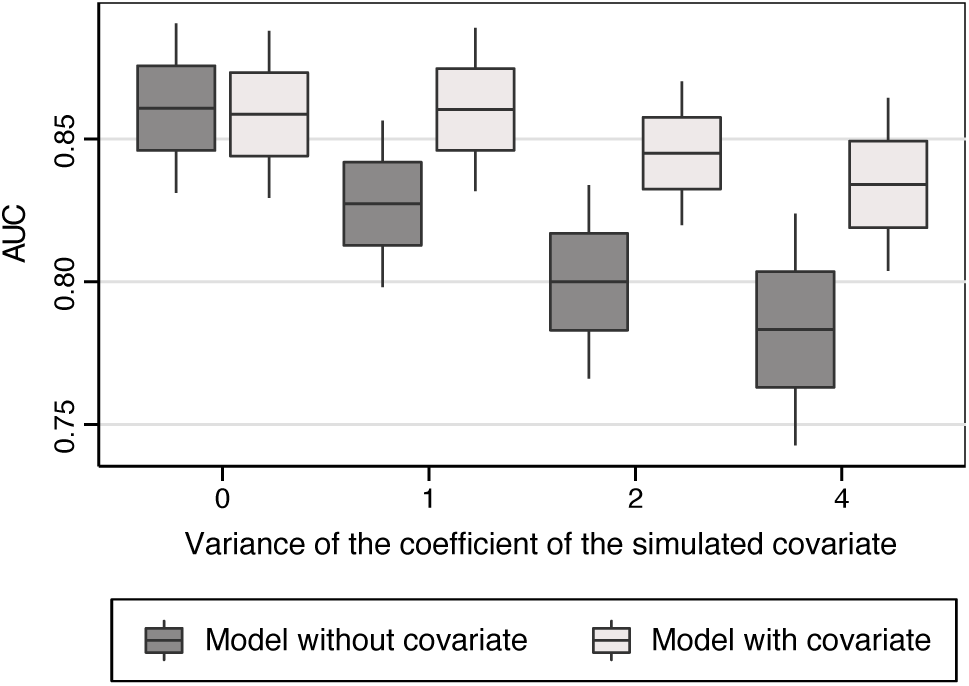
The network recovery ability (AUC) as a function of the confounding covariate strength *γ* for the agnostic MAGMA method vs. the MAGMA method with covariate effect. We considered 20 simulation iterations for networks of size 200 with an average degree of 2, number of samples *n* = 100, conditioning number *k* = 10, graph structure was random, read count data were simulated according to a ZINB.

#### 3.1.7 Comparison with other association network approaches

We compare the existing methods regarding the presence or absence of structure among samples due to a covariate. As shown in Figure 4, MAGMA showed better performances than the three reference methods in reconstructing microbial interactions, namely SparCC, CoNet and SPIEC-EASI. The networks recovered by CoNet are derived from the calculation of Spearman correlation p-values by permutation and bootstrap. In our simulations, this did not have an added value compared to the networks obtained from Spearman correlations thresholding. The Pearson correlation network and the graphical lasso model on raw data did not work well without data normalization: linear correlations should not be calculated from raw read count data. The graphical lasso with nonparanormal SKEPTIC transformation had a higher AUC than that obtained with SparCC and SPIEC-EASI; yet this non-parametric transformation is typically not used for the study of microbiota data. In the presence of a covariate, the performance of all competing methods degraded significantly and the AUC dropped. Under our simulations, MAGMA inference yielded the best performance. We therefore conclude that it is essential to take into account the potential covariates with structural effects on the microbiota.

**Figure 4:**
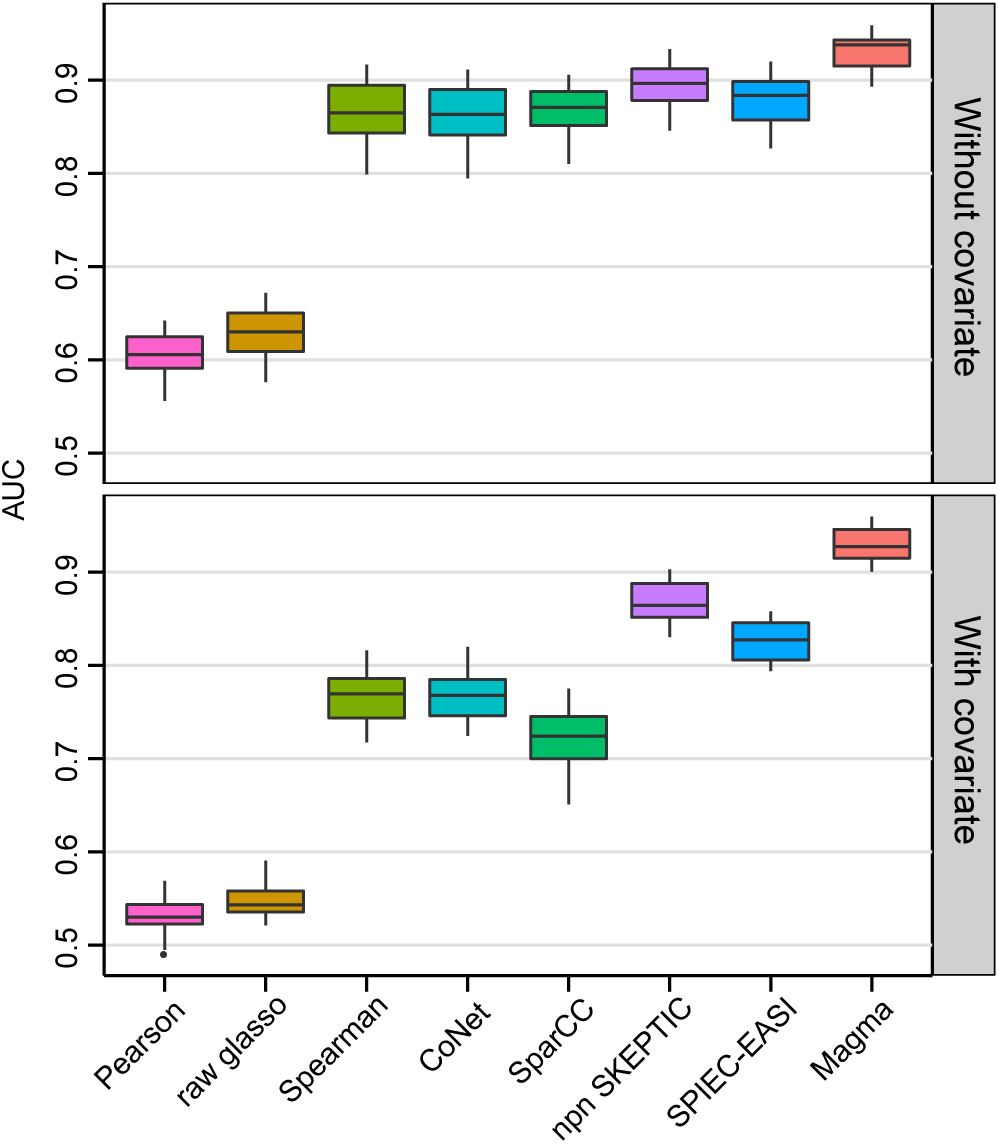
Comparison of inference methods considering a covariate effect. Boxplots of the AUC criterion for different network inference methods. We considered 20 simulation iterations for networks of size 200 with an average degree of 2, number of samples *n* = 300, condition number *k* = 5, graph structure was random and read count data were simulated according to a ZINB. The quantitative covariate parameter *γ* was drawn from a *N* (0, 2), when covariate was effective. CoNet designates network obtained from Spearman correlation p-values by 100 iterations of permutation and bootstrap. For SparCC, the correlation threshold parameter was equal to 0.3 and 100 iterations were done in the outer loop and 20 in the inner loop. raw Glasso, npn SKEPTIC, SPIEC-EASI, and MAGMA were network obtained by graphical lasso inference from raw data, nonparanormal SKEPTIC transformation, clr transformation and MAGMA transformation respectively. For Pearson, Spearman and SparCC networks, we computed the path of inferred networks by thresholding the correlations. For CoNet, we thresholded the p-values. For the graphical lasso, we varied the regularization parameter.

### 3.2 Microbial data illustration: Human Microbiome Project

In this section, we present the analysis of the 16S variable region V3-5 data from the Human Microbiome Project (HMP) [4, 5]. The study collected microbiomes of healthy individual at various body sites. The data was retrieved on the qiita data platform [58, study ID 1928]. This study brings together a total of 6,000 samples from 18 different the human body sites. A total of 10,000 microbial species occupy the human ecosystem. We first studied a stool microbiota sample, comparing MAGMA with SparCC and SPIEC-EASI. Second, we analyzed the stool and saliva microbiota in a single study in order to show the usefulness of the covariate implementation in the MAGMA method.

#### 3.2.1 Inference of gut microbiota network

Among other roles, the gut microbiota is involved in nutrient metabolism and in the prevention of colonization by pathogenic micro-organisms. Getting insight into the functioning of this microbial ecosystem is therefore a critical scientific issue. Stool HMP data contains 388 samples and 10,730 OTUs, with most OTUs being rare. We filter out OTUs present in less than 25% of the samples and remove the samples whose sequencing depth is less than 500 reads on the remaining OTUs. These samples show large stochastic variability and in a properly weighted analysis would not add much information. After this preprocessing we obtain an OTU table with 360 samples and 306 OTUs.

Figure 5A show the stool network obtained by MAGMA, SparCC and SPIEC-EASI. With the *stars* selection from the `huge` R package, MAGMA and SPIEC-EASI selected a little over 2000 edges (2356 for MAGMA, 2332 for SPIEC-EASI). For comparative purposes each network is shown with the same amount of 2000 edges. Figure 5B shows the network node degree distributions as well as the Venn diagrams of the inferred links. SparCC network has the wides distribution with high degree nodes for both positive and negative association links. Regarding positive links, the SPIEC-EASI network has more nodes characterized by low degrees than the other methods.

**Figure 5:**
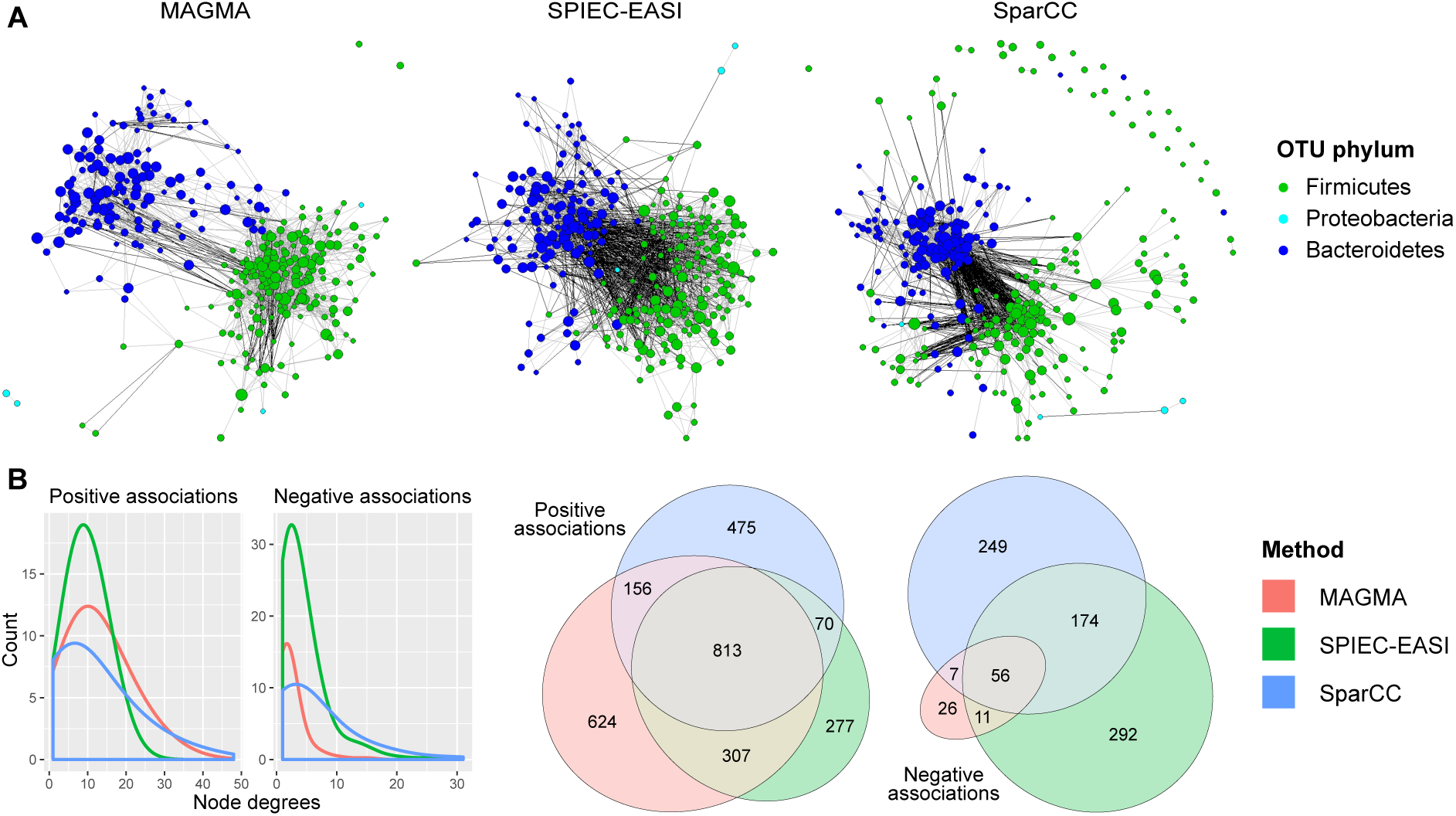
Stool microbiota network. (**A**) Stool microbial association network obtained from three methods MAGMA, SPIEC-EASI and SparCC. Nodes are OTUs. Black and gray links represent negative and positive associations, respectively. (**B**) positive and negative associations obtained from the three networks: smoothed histograms of node degrees and Venn diagrams representing the overlap of inferred links between the different methods.

The three networks show a strong antagonism between the groups of the Firmicutes and Bacteroidetes phila. MAGMA network showed the most tempered opposition between this two groups and has fewer negative links (100) than the other networks (486 for SparCC and 533 for SPIEC-EASI). Less than half of the negative links recovered by SparCC and SPIEC-EASI were identical, raising questions about their veracity. The MAGMA stool network has more positive links: 25% and 30% more than SparCC and SPIEC-EASI respectively. Relative to this, only 33% of positive links recovered by MAGMA differed from those found by these two methods. Compared to other tools, MAGMA seems to identify a coherent network with sensible biological structure, and it showed a good reproducibility of results compared to the other methods.

#### 3.2.2 Microbial network body site variation

To illustrate MAGMA’s ability to account for confounding or, put differently, to analyze information across heterogeneous samples, we pool two different sets of HMP microbiota and introduce a covariate. We group gut microbiota data and salivary microbiota data and introduce a, probably marked, “body site” effect. Again, we filter out OTUs present in less than 25% of the samples and remove samples with sequencing depth of less than 500 reads on the remaining OTUs. This results in an OTU table with 665 samples and 245 OTUs.

Figure 6 shows the two networks we obtain with and without integration of the body site covariate in MAGMA. In the network *without* covariate (Figure 6A), two sets of OTUs were stand out. A first group with Firmicutes and Bacteroidetes phyla corresponds to intestinal microbiota and a second group corresponds to salivary microbiota. The OTUs of the same group are positively associated with each other, while two OTUs of different groups are negatively related. In the network *with* covariate (Figure 6B), there were again two groups of OTUs. This time the spurious negative links between the two groups disappear, because the difference in frequency of OTUs between body sites has been taken into account by means of the body site covariate, which allows to find real functional interactions between the various OTUs. This includes various positive associations between the two groups of OTUs. In fact, there are no common negative links between network A and B. Negative correlations due to the average body site effect are shifted to 0 when normalizing by considering the covariate (Figure 6C). Positive correlations due to OTU co-presence in a specific microbiota are centered when including the covariate. Taking into account the body site in MAGMA makes it possible to obtain a *consensual* network.

**Figure 6:**
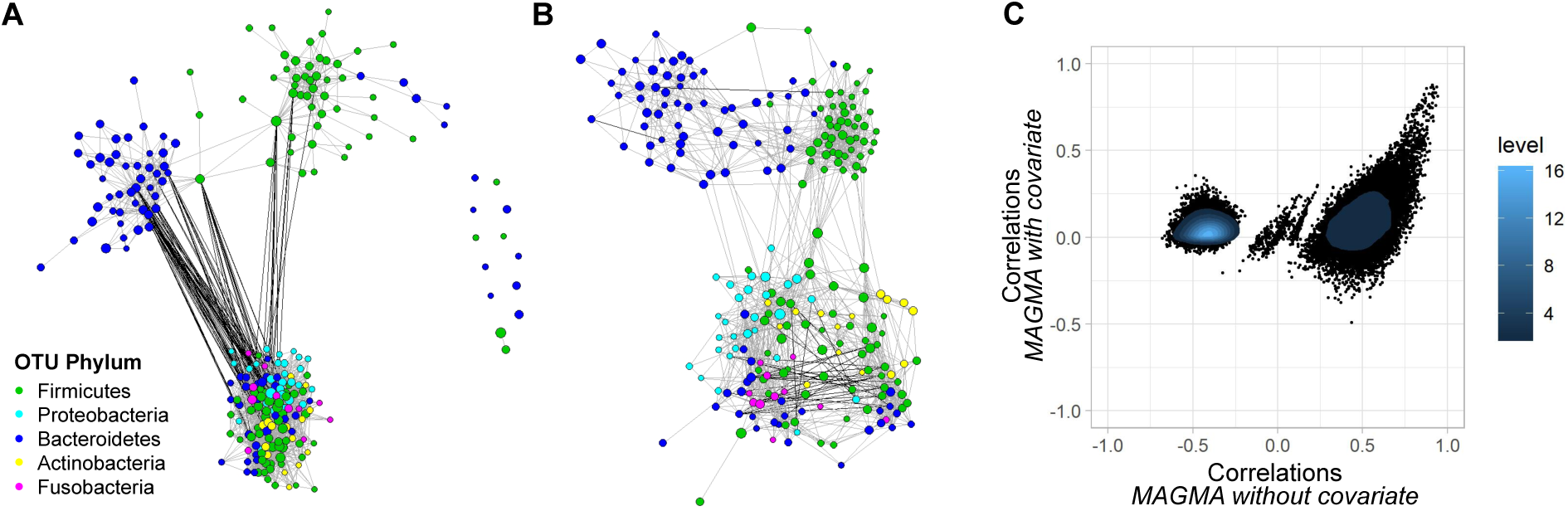
Association network of stool and saliva microbiota pooled data. (**A**) Stool and saliva microbial association network without body site covariate. (**B**) Stool and saliva microbial association network including body site factor. Regularization parameters for the two networks were determined with stars selection. (**C**) Correlations of MAGMA without including body site covariate versus correlations of MAGMA including body site effect. Correlations were computed from Pearson correlations of MAGMA transformed data (normalization defined by equation (7)).

## 4 Conclusion

We have introduced a network model that responds to the methodological challenges arising from sequencing read count data: excess of zeros, over-dispersion, compositionality and the presence of covariates. To meet these challenges, the network inference method we propose takes advantage of a GLM-inspired parametric mapping function, while being based on the well-known Gaussian graphical model. MAGMA offers a normalization approach based on the theory of copulas. Moreover, it takes into account variable sequencing depth estimating the library size effect by the geometric mean of pairwise ratios.

In the simulation studies we show that the approximations made during the transformation of the data rapidly converge towards the correct solution when the number of samples and the strength of the correlations increase. The ZINB law we propose is flexible and can deal easily with moderate amount of model misspecification. The integration of covariates improves the quality of the inference in presence of structural factors affecting the OTU read counts. Finally, MAGMA performs better than other available competitors in a wide variety of situations.

We have applied MAGMA to infer an intestinal microbial network from a HMP data, allowing for heterogeneous samples. The resulting network shows a consensual interactions that are not affected by wildly different OTU counts for various body sites. All this shows that MAGMA is a practical tool for inferring microbial functional networks from metagenomic sequence read count data.

## Acknowledgments

The work was funded by two INRA metaprogrammes: Meta-omics of microbial ecosystems (MEM) and Integrated management of animal health (GISA) and by the European Cooperation for Statistics of Network Data Science (COSTNET: COST Action CA15109).

